# How antisense transcripts can evolve to encode novel proteins

**DOI:** 10.1101/2023.08.30.555508

**Authors:** Bharat Ravi Iyengar, Anna Grandchamp, Erich Bornberg-Bauer

**Affiliations:** Institute for Evolution and Biodiversity, University of Mü nster, Hü fferstrasse 1, 48149 Mü nster, Germany; Department of Protein Evolution, Max Planck Institute for Biology Tü bingen, Max-Planck-Ring 5, 72076 Tü bingen, Germany

## Abstract

Protein coding features can emerge *de novo* in non coding transcripts, resulting in emergence of new protein coding genes. Studies across many species show that a large fraction large fraction of evolutionarily novel non-coding RNAs have an antisense overlap with protein coding genes. The open reading frames (ORFs) in these antisense RNAs could also overlap with existing ORFs. In this study, we investigate how the evolution an ORF could be constrained by its overlap with an existing ORF in three different reading frames. Using a combination of mathematical modeling and genome/transcriptome data analysis in two different model organisms, we show that antisense overlap can increase the likelihood of ORF emergence and reduce the likelihood of ORF loss, especially in one of the three reading frames. In addition to rationalising the repeatedly reported prevalence of *de novo* emerged genes in antisense transcripts, our work also provides a generic modeling and an analytical framework that can be used to understand evolution of antisense genes.

## Introduction

New protein coding genes often arise from existing protein coding genes. This process frequently involves duplication of an existing gene, and a subsequent divergence of one of the duplicated copies from the ancestral sequence (Long *et al*., 2003; Rastogi and Liberles, 2005; Näsvall *et al*., 2012). Several studies have shown that protein coding genes can also emerge *de novo*, in DNA sequences that did not previously encode a protein (*de novo* gene emergence; Tautz and Domazet-Lošo, 2011; Zhao *et al*., 2014; Schmitz and Bornberg-Bauer, 2017; Vakirlis *et al*., 2017; Van Oss and Carvunis, 2019; Vakirlis *et al*., 2020). A protein coding gene thus emerged does not inherit the DNA sequence features necessary for gene expression (transcription and translation), from an ancestral protein coding gene. It must therefore acquire them through random mutations.

Among the most fundamental features necessary for transcription, are a promoter and a poly-adenylation signal (polyA signal). Promoters help to initiate transcription by recruiting RNA polymerase (Lenhard *et al*., 2012; Haberle and Stark, 2018). However, transcription in eukaryotes can be initiated at a low rate even in the absence of a defined promoter sequence (Clark *et al*., 2011). A polyA signal not only marks the end of transcription but also elicits an enzymatic addition of a polyadenosine tail (polyA tail) to the 3’ end of the transcribed RNA. The polyA tail makes the RNA resistant to degradation, and facilitates its export to cytoplasm where it can be translated by the ribosome (Richard and Manley, 2009; Proudfoot, 2011; Stewart, 2019). The most basic requirement for translation is an open reading frame (ORF), which is the region of an RNA that is translated into a protein sequence. Efficient translation often requires additional features such as Kozak consensus sequences (Kozak, 1986; Acevedo *et al*., 2018; Noderer *et al*., 2014), an optimal codon usage (Hanson and Coller, 2017), and other context dependent regulatory features present in the 5’ and 3’ untranslated regions of the RNA (Hinnebusch *et al*., 2016; Mayr, 2017).

Because heritable (germline) mutations are rare in most organisms (less than 1 mutation in 100 million base pairs of DNA per generation; Schrider *et al*., 2013; Zhu *et al*., 2014; Jee *et al*., 2016), it is unlikely for many features to emerge simultaneously. That is, features must evolve sequentially. This in turn means that emergence of a phenotype, such as gene expression, is more likely when some required features already exist, and the missing features emerge via mutations. For example, *de novo* emergence is more likely when an ORF is already present and transcriptional features emerge subsequently, or *vice versa*. In our recent work, we also show that *de novo* emergence is more likely via the trajectory where transcription emerges before the emergence of an ORF (Iyengar and Bornberg-Bauer, 2023). Thus stably synthesized RNAs that are not actively and specifically involved in protein synthesis (such as long non-coding RNAs or lncRNAs) can be good sources of new proteins.

Experimental analyses of the ribosome’s footprint on RNAs (ribosome profiling) suggest that some ORFs present in lncRNAs are actively translated (Ruiz-Orera *et al*., 2014; Ingolia *et al*., 2014; Patraquim *et al*., 2022; Blevins *et al*., 2021; Wacholder *et al*., 2023). Proteins synthesized from the translation of such ORFs can also be beneficial to the host organism (Patraquim *et al*., 2022; Wacholder *et al*., 2023). Many lncRNA genes share their genomic location with other genes, but are transcribed in the opposite direction (antisense overlap; Mattick *et al*., 2023). A recent study has characterized previously unknown RNAs in different species of yeasts, and has shown that a large proportion of these RNA genes have an antisense overlap with existing genes (Blevins *et al*., 2021). This study also shows that ORFs contained in these RNAs show signatures of translation. These translated ORFs also include those that have recently emerged in one specific species of yeast. However, these species specific ORFs are less efficiently translated than the ORFs that are conserved between different species. Overall, this study lends support to a hypothesis that many new proteins arise from antisense RNAs. It is likely that the ORFs encoding such proteins are also antisense to existing genes.

In this study, we analyse the emergence of ORFs in antisense RNAs. We specifically focus on ORFs that have an antisense overlap with the coding region (canonical ORF) of an existing protein coding gene. We refer to these ORFs as antisense ORFs (asORFs). Evolution of asORFs is also interesting because it is constrained by the evolutionary selection pressure on the overlapping protein coding genes (Sabath *et al*., 2012; Mir and Schober, 2014). A pair of mutually antisense ORFs can overlap with each other in three different reading frames. That is, the codon positions in the two ORFs can either perfectly overlap or be offset by one or two nucleotides. The constraints on the co-evolution of the two ORFs would be different in the different reading frames (Mir and Schober, 2014). Our study aims to explore the constraints that affect the evolution of asORFs. To this end, we employ a mathematical model to calculate the probabilities of asORF emergence and loss, in each of the three reading frames. Using the model, we predict that one of the reading frames has a higher propensity to harbor ORFs. We also predict that the likelihood of ORF emergence in this reading frame is higher, and that of ORF loss is lower, than in the other two reading frames. We support our model’s predictions with genome analysis of two different organisms – *Saccharomyces cerevisiae* and *Drosophila melanogaster*. We also find that emergence of asORFs in reading frame 1 can be more likely than emergence of non-antisense (intergenic) ORFs.

## Results

We developed a mathematical model to estimate the probabilities of ORF emergence and loss, in DNA regions antisense to existing protein coding ORFs. This model is defined by two kinds of probability. The first is the probability of finding a certain kind of DNA sequence, for example an ORF. This stationary probability depends on the nucleotide composition of the DNA region that can be roughly approximated by GC-content or by the frequencies of short DNA sequences (oligomers). The second kind of probability describes the mutational change of a sequence to a different kind of sequence. For example, gain or loss of an ORF. This transition probability depends on the mutation rate and mutation bias, in addition to nucleotide composition. We estimate these parameters primarily from the data on the yeast, *Saccharomyces cerevisiae* (Table 1; Zhu *et al*., 2014). Our choice is motivated by the fact that the budding yeast is a convenient model organism for laboratory experimental studies that can be used to validate several of our theoretical predictions. We also performed analogous analyses using data obtained from *Drosophila melanogaster* (Schrider *et al*., 2013, Supplementary Material).

**Table 1:** Mutation bias probabilities for different nucleotide mutations in *Saccharomyces cere- visiae* (Zhu *et al*., 2014). `A:T` denotes an `A`-`T` base pair in a double stranded DNA. Thus `A`*→*`G` mutation on one DNA strand would cause a `T`*→*`C` mutation on the complementary strand. We describe the other mutations in the same way. For our model, we used the reported mutation rate of 1.7 *×* 10*^−^*^10^ mutations per nucleotide position per generation, in diploid *Saccharomyces cerevisiae* cells (Zhu *et al*., 2014). For mutation bias probabilities in *D. melanogaster*, see Table S1.

| Substitution | Probability($\mu$) |
| --- | --- |
| A : T → T : A | 0.063 |
| A : T → G : C | 0.144 |
| A : T → C : G | 0.110 |
| G : C → A : T | 0.182 |
| G : C → T : A | 0.349 |
| G : C → C : G | 0.152 |

We estimated the stationary and transition probabilities of antisense ORFs (asORFs, Equations 1 – 3) using the existing (sense) ORF as a reference. asORFs can overlap with the sense ORFs in three different reading frames (henceforth referred to as just “frames”). In frame 0, the codons in the asORF exactly overlap the codons in the sense ORF. In frames 1 and 2, the codons in the asORF are shifted towards the 5’ end of the sense ORF by one and two nucleotide positions, respectively. Thus in frames 1 and 2, the sequence of an antisense codon is determined by two partially overlapping sense codons (dicodons, Figure 1A). Due to this sequence overlap, the evolution of asORFs would be constrained by the evolutionary selection pressures on the sense ORF. Furthermore, these constraints would be different for asORFs located in the three different frames. We analysed the evolution of asORFs when the sense ORF is under three different levels of purifying selection. The first level describes an absence of purifying selection, where any kind of mutation except a non-sense mutation (gain of stop codon) in the sense ORF is tolerated. The second level describes a weak purifying selection that allows synonymous mutations, as well as mutations where an amino acid is substituted by a chemically similar amino acid (for example, aspartic acid to glutamic acid; Table 4). Finally, the third level describes a strong purifying selection, where only synonymous mutations are tolerated in the sense ORF.

**Figure 1:**
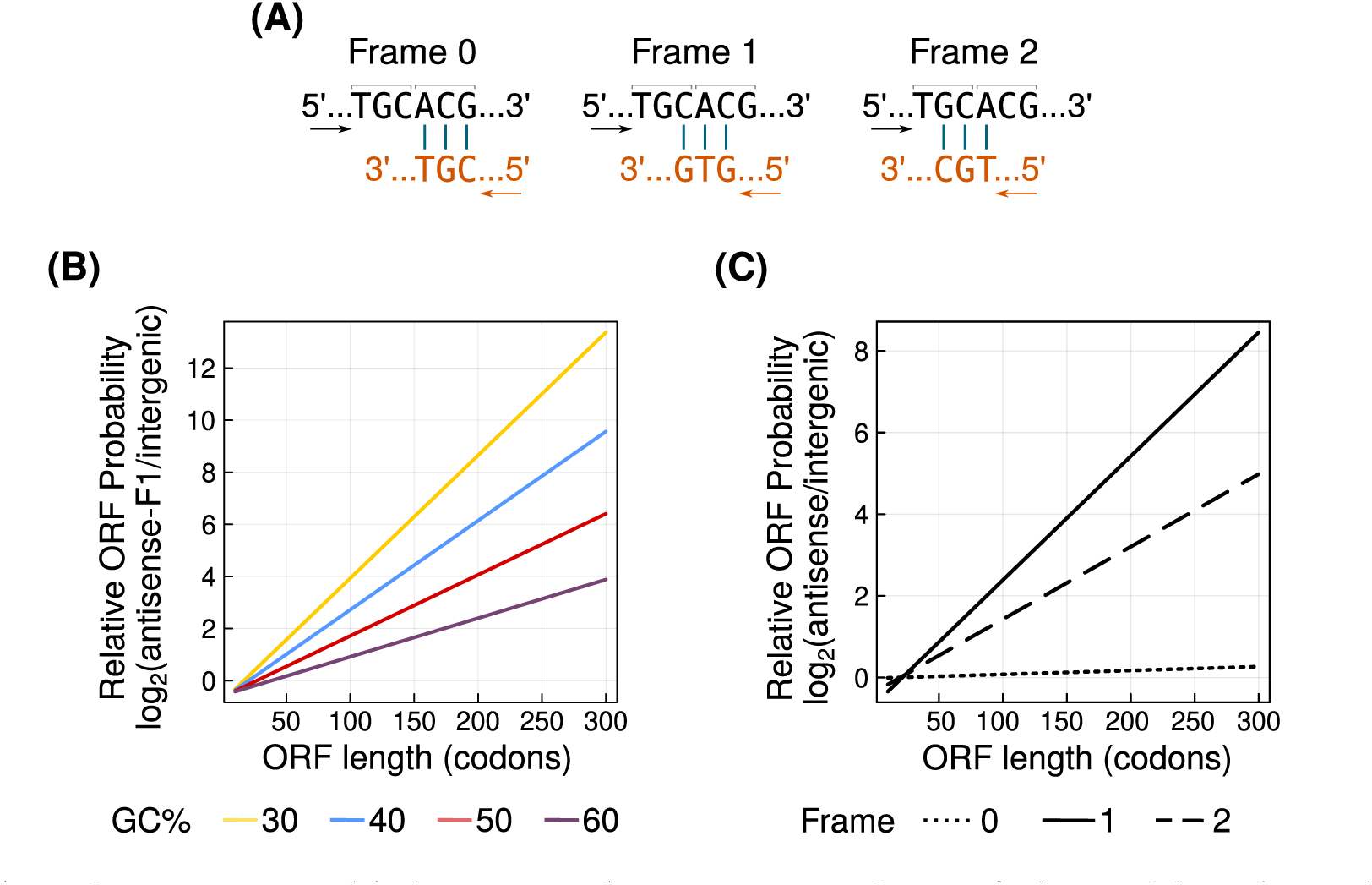
asORFs are more likely to exist than intergenic ORFs of identical lengths and composition. **(A)** A hypothetical antisense codon (bottom sequence, orange) can overlap with sense ORF (top sequence, black) in three different frames. Arrows indicate the direction of translation and vertical bars indicate base complementarity. Adjacent codons in the sense ORF are demarcated with horizontal square brackets. **(B)** The probability of asORFs in frame 1 relative to that of intergenic ORFs (log_2_ ratio, vertical axis). Line colors indicate the GC-content of the ORFs (yellow = 30%, blue = 40%, red = 50%, purple = 60%). We do not show asORFs in frames 0 and 2 because their probabilities are identical to that of intergenic ORFs. **(C)** The probability of asORFs relative to that of intergenic ORFs (log_2_ ratio, vertical axis), calculated using frequencies of short DNA sequences from the yeast genome. Frames 0, 1 and 2 are denoted by dotted, solid and dashed lines, respectively. Horizontal axes in panels **(B)** and **(C)** show the length of the ORFs. We only show asORFs that overlap completely with the sense ORF.

### Antisense ORFs are more likely to exist in frame 1

For any stretch of DNA to be an ORF, its sequence should contain 3*n* nucleotides (*n ≥* 3), with a start codon that marks its beginning, and exactly one stop codon that marks its end. The absence of any stop codon within the DNA sequence is the most important factor in determining the existence of an ORF. That is because the likelihood of a premature stop codon increases exponentially with the ORF’s length, whereas the likelihoods of a start codon and a terminal stop codon are independent of the ORF’s length (Equations 1 – 3).

Based on these considerations, we determined the probability of finding an asORF. To this end, we first calculated the probability of finding a stop codon in the three antisense frames. A stop codon can exist in frame 0 wherever the three reverse complementary codons exist in the sense ORF. It can exist in frames 1 and 2, overlapping with one of the 128 and 192 dicodons in the sense ORF, respectively. We calculated the probability of finding (or not finding) an antisense stop codon based on the probability of finding the corresponding sense codons and dicodons. Codon and dicodon probabilities depend on the nucleotide composition, which can be approximated by the GC-content of the locus (Iyengar and Bornberg-Bauer, 2023). We calculated the probability of a start codon without considering the effect of antisense overlap because this effect would be small in magnitude. Using the start and stop codon probabilities, we estimated the probability of finding an asORF of different lengths in each of the three frames. We did so for four different values of GC-content (30, 40, 50 and 60%). We found that frame 1 has the highest likelihood of harboring an asORF in most cases. The only exceptions are ORFs shorter than 39 codons present in a DNA region with a GC-content of 60%. Even for these exceptional cases, the probability of an asORF in frame 1 is no less than 91% of the corresponding ORF probabilities in the other frames.

To understand whether asORFs are more likely to be found than ORFs in intergenic regions, we compared the probabilities of asORFs with that of intergenic ORFs of identical GC-content and lengths. We note that that the probability of intergenic ORFs is not constrained by existing ORFs and is thus purely dependent on nucleotide composition. We found that the probability of asORFs in frames 0 and 2 were identical to that of the corresponding intergenic ORFs, which means that asORFs in frame 1 are more likely to exist than intergenic ORFs except in the few cases where the ORFs have a GC-content of 60% and less than 39 codons (Figure 1B). We expect that intergenic ORFs can indeed be more numerous than asORFs if intergenic regions are long. Our results merely suggest that given that length and GC-content are identical, the probability of an ORF increases when it has an antisense overlap with an existing ORF in frame 1.

We also calculated the probability of asORFs using actual codon and dicodon frequencies in annotated yeast ORFs. Likewise, we calculated the probability of intergenic ORFs using the frequencies of DNA trimers in yeast intergenic genome. With this analysis, we found that asORFs longer than 54 and 75 codons in frames 1 and 2, respectively, are more likely to exist than intergenic ORF of the same lengths (Figure 1C). For all ORF lengths, asORFs in frame 0 are less likely to exist than intergenic ORFs.

Our supplementary analyses using *D. melanogaster* as a model for mutation bias produced similar results (Figure S1A). However, when we computed the ORF probabilities using the frequencies of codons, dicodons and intergenic trimers from *D. melanogaster*, we found that frame 0 was most likely to harbor an asORF (Figure S1B). This small deviation from a GC-content based prediction could result from a biased codon usage in *D. melanogaster* protein coding genes.

### Antisense ORFs are frequently located in frame 1

Our mathematical model predicts that frame 1 is more likely to harbor asORFs than the other two frames. To verify this prediction, we analysed the genome of the budding yeast, *S. cerevisiae*. We specifically chose this yeast as a model because most of its genes lack introns. This in turn allows us to investigate asORFs whose overlap with the sense ORFs is not interrupted by intronic sequences. Our choice of yeast as a model was further motivated by the availability of data on novel antisense RNAs identified in a recently published study (Blevins *et al*., 2021). This study further showed that new protein coding genes can emerge *de novo* from these antisense RNAs. We identified all asORFs located in the novel RNAs reported in this study, and calculated the frame in which they overlap with the annotated (sense) ORFs. We also included seven annotated yeast antisense RNAs for the identification of asORFs. Next, we calculated the number of asORFs in each of the three frames, that are at least 30nt long and are wholly contained within the boundaries of a sense ORF. We found that *∼*39% of all asORFs were located in frame 1, while *∼*33% and *∼*28% asORFs were located in frames 2 and 0, respectively (Table 2, Figure 2). We also calculated the number of ORFs that have at least 50% of their sequences overlapping in antisense with a sense ORF. This relaxation of overlap percentage did not remarkably increase the number of identified asORFs. To understand if the observed number and proportion of asORFs are in agreement with the model, we calculated the expected number of asORFs in each frame (Equation 4). Specifically, we estimated the total number of expected ORFs that are at least 30nt long and are located in genomic region where antisense RNAs overlap with a known ORF.

**Table 2:** Expected and observed numbers of antisense and intergenic ORFs. For both expected and observed number of asORFs, we only consider ORFs that overlap completely with a sense ORF. Here “sub-ORF” refers to smaller ORFs that exist within an ORF such that they begin at any position where a methionine is encoded, and end at the stop codon of the long ORF. We do not include sub-ORFs shorter than 30nt.

|  | Total loci | Expected ORFs | Observed ORFs | Observed + sub-ORFs |
| --- | --- | --- | --- | --- |
| Antisense F0 | 7921647 | 626 | 447 | 494 |
| Antisense F1 | 7921647 | 651 | 646 | 903 |
| Antisense F2 | 7921647 | 587 | 548 | 623 |
| Intergenic | 1635429374 | 49785 | 40647 | 48598 |

**Figure 2:**
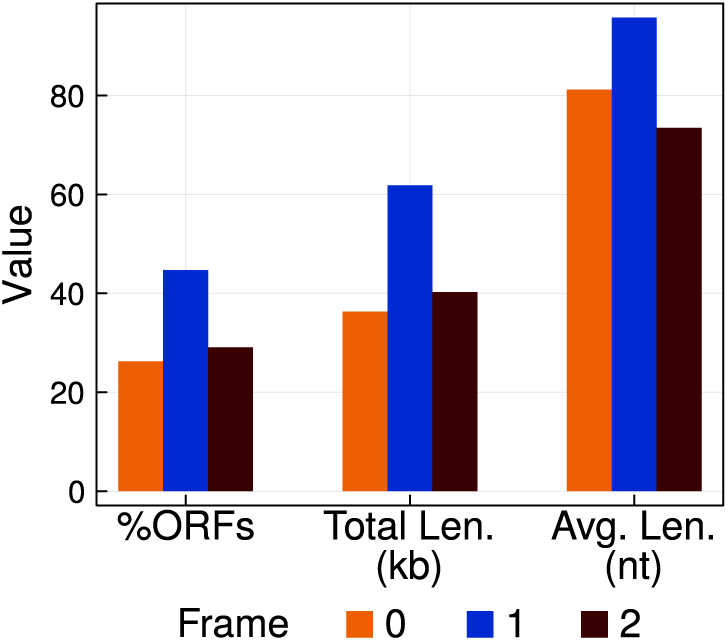
Yeast asORFs are preferentially exist in frame 1 than in the other two frames. We show three metrics of frame preference as three bar groups – percentage of total ORFs (left), total (combined) ORF length (middle), and average ORF length (right), in each of the three frames (bar colors – 0: orange, 1: blue, and 2: brown). We do not show ORFs that overlap partially with the sense ORF.

We found that the actual asORFs in the yeast genome were 0.7 – 28% fewer than expected (Table 2). The ORF identification tool we used (*getorf* ; Rice *et al*., 2000), reports the longest ORF. However, alternate start codons can exist within the ORF sequence wherever a methionine is encoded. Our model does not reject short ORFs (sub-ORFs) within a longer ORFs. When we included the sub-ORFs (*≥*30nt), the observed number of asORFs in frame 1 was significantly larger than its expected number (one-tailed Fisher exact test, *P <* 10*^−^*^15^; Table 2). We further note that our calculation of expected number of asORFs (Equation 4) assumes that existence of ORFs in the three different frames is independent of each other. However, presence of an ORF in any one frame can reduce the probability of ORFs in overlapping alternate frames.

Probability of finding an ORF can not only determine the expected number of ORFs, but also the expected length of ORFs. Therefore, we next asked if asORFs in frame 1 are longer than those in the other two frames. To this end, we calculated the average ORF length, and the combined length of all ORFs, in each frame. We found that asORFs in frame 1 were on an average 96nt long, whereas asORFs in frames 2 and 0 were on an average 81nt and 72nt long, respectively (Figure 2). Furthermore, the total (cumulative) length of all the asORFs in frame 1 was higher than that of the asORFs present in the other two frames. Specifically, the combined ORF length in frames 0, 1 and 2 was 36kb (kilobases), 62kb and 40kb, respectively (Figure 2).

Next, we analysed if the observed frequency of intergenic ORFs is different from that of asORFs. To this end, we calculated the observed number of intergenic ORFs including the sub-ORFs, in *S. cerevisiae* genome, using a procedure identical to that we used for identifying asORFs. We then compared the frequencies of intergenic ORFs (observed ORFs relative to total loci, Table 2) with that of each type of asORFs, and found that the frequencies of all the three types of asORFs were higher than that of intergenic ORFs (one-tailed Fisher exact test, *P <* 10*^−^*^15^). We note again that this result does not indicate that intergenic ORFs are less likely to occur than asORFs, as we show that they are indeed more numerous than asORFs (Table 2).

We also performed a similar analysis of *D. melanogaster* genome. Specifically, we used genome and transcriptome data from inbred lines obtained from seven geographically distinct *D. melanogaster* populations (Grandchamp *et al*., 2023). We used these datasets because they contain several novel RNAs that are not annotated in the reference genome. We found that among the three antisense frames, frame 1 harbored the most number of asORFs. The cumulative length of all the asORFs in the frame 1 was also higher than those in the other two frames (Figure S2). This was true for all the seven lines, and also for the set of unique orthologous sequences between all the lines (orthogroups). However, asORFs in frame 1 were not always, on an average, longer than those in the other two frames. Specifically in five out of seven lines, frame 0 contained on an average the longest ORFs. This was also the case for the sequences in the orthogroups. However, in these lines (as well as the orthogroups) the average length of asORF in frame 1 was no less than 86% of the average length of asORF in frame 0. A possible reason for the larger average length of asORF in frame 0 could be the codon usage bias in *D. melanogaster* protein coding genes (Vicario *et al*., 2007). We also analysed if intergenic ORFs have a higher frequency than asORFs in *D. melanogaster*. We restricted this analysis to asORFs that completely overlap with a coding exon, and found that intergenic ORFs have significantly higher frequency than asORFs (one-tailed Fisher exact test, *P <* 0.01). This deviation from the prediction could result from our highly stringent analysis of asORFs, and due to the existence of several pseudogenes and transposable elements in the annotated intergenic genome.

Overall, our genome data analyses from both organisms support our model’s prediction that frame 1 offers the most optimal location for asORFs.

### Antisense overlap can facilitate ORF emergence and reduce ORF loss

We next analysed how likely it is for asORFs to emerge, when they are not already present. To this end, we calculated gain probability of asORFs in each of the three frames, and under three different intensities of purifying selection. We also calculated the probability of ORF gain in the intergenic regions. We found that asORFs are less likely to emerge in frames 0 and 2, than ORFs in intergenic regions, for all ORF lengths and GC-content. In contrast, the emergence of asORF emergence in frame 1 is often more likely than that of intergenic ORFs, especially when the ORFs are long (Figure 3A).

**Figure 3:**
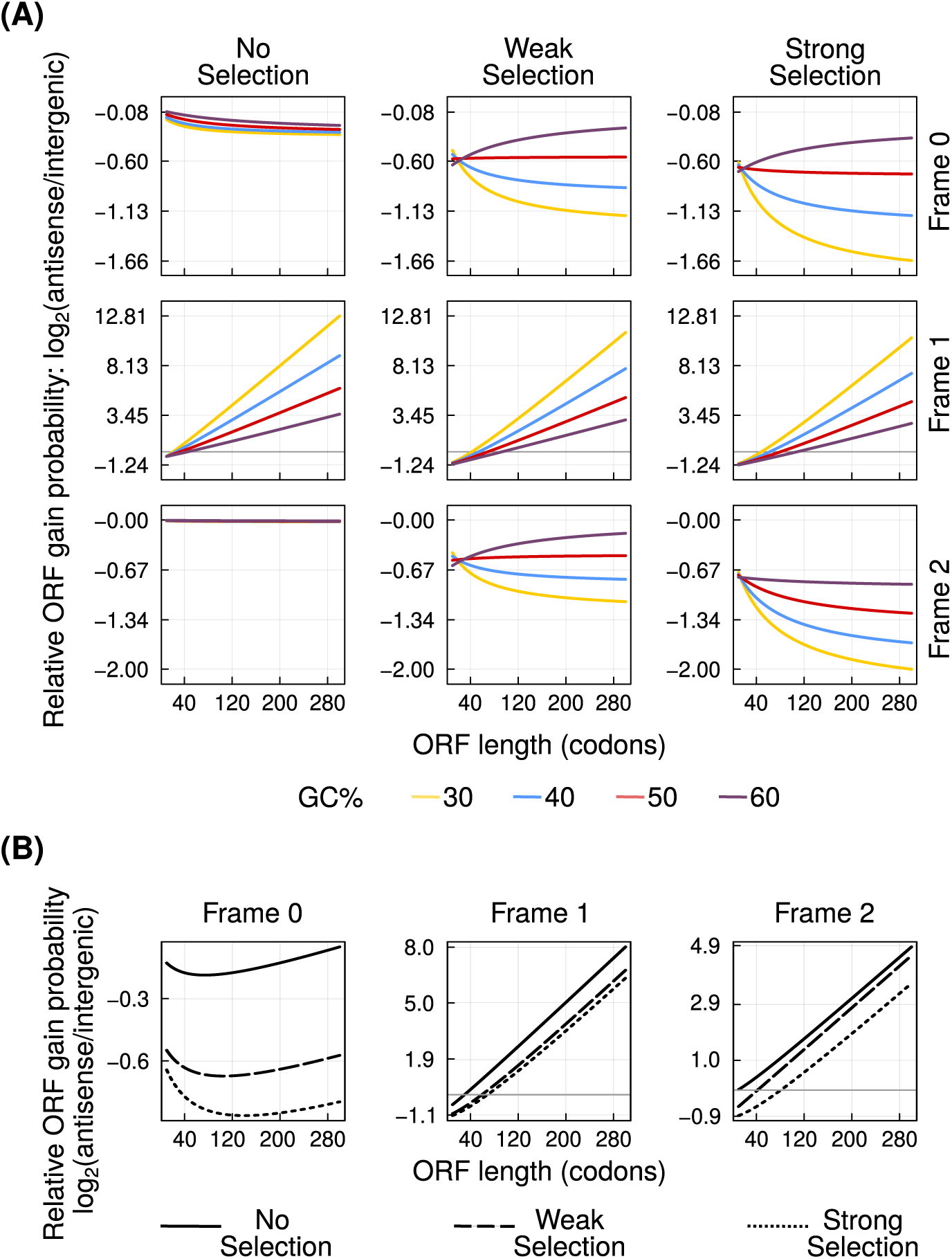
Antisense overlap can facilitate ORF emergence. **(A)** The probability of ORF emergence in the three antisense frames (left to right) relative to that in intergenic regions (log_2_ ratio, vertical axis), at different intensities of purifying selection (top to bottom). Line colors indicate the GC-content of the ORFs. **(B)** ORFs gain probability in the three antisense frames relative to that in intergenic regions (log_2_ ratio, vertical axis), calculated using frequencies of short DNA sequences from the yeast genome. Dotted, solid and dashed lines, denote the zero, weak and strong purifying selection, respectively. Horizontal axis in every plot shows the length of the ORFs. In every plot, we only show asORFs that overlap completely with the sense ORF.

Increasing the intensity of purifying selection reduces the emergence likelihood of asORFs in all the three frames. However, emergence of asORFs in frame 1 is still more likely than emergence of intergenic ORFs, even under strong purifying selection (Figure 3A). Specifically, the minimum ORF length where asORFs in frame 1 are more likely to emerge than intergenic ORFs, increases with GC-content and the intensity of selection. For example, in the absence of purifying selection, and at a GC-content of 30%, this length is smaller than the smallest investigated ORF length of 30 codons. At the same intensity of selection, this length is 46 codons when the GC-content is 60%. ORFs can emerge in antisense frame 1 more frequently than in intergenic regions under strong purifying selection, with GC-contents of 30% and 60%, if they are longer than 49 and 109 codons, respectively.

Our analysis of ORF gain probabilities using the frequencies of DNA oligomers (codons, dicodons and intergenic trimers), also shows that asORFs are very likely to emerge in frame 1 (Figure 3B). ORFs longer than 30 (shortest investigated length), 60 and 69 codons are more likely to emerge in antisense frame 1 than in intergenic regions, when the purifying selection is absent, weak and strong, respectively. Interestingly, this analysis revealed that, although asORFs are less likely to emerge in frame 2 than in frame 1, they can emerge more frequently than intergenic ORFs. Specifically when the purifying selection is absent, weak and strong, ORFs that are more likely to emerge in antisense frame 2 than in intergenic regions, contain at least 30, 43 and 82 codons, respectively.

Our analogous analysis with parameters estimated from *D. melanogaster* also produced similar results (Figures S3).

Purifying selection reduces the number of tolerated mutations in a DNA locus. We note again that even the lowest intensity of purifying selection according to our definition, disallows nonsense mutations from occurring in the sense ORFs. We thus hypothesized that overlap with a sense ORF may protect the asORFs from being lost. To this end, we calculated ORF loss probabilities for different ORF lengths, GC-content, and intensities of purifying selection (Figure 4A). In an analogous analysis, we used codon, dicodon, and intergenic trimer frequencies, instead of GC-content, to calculate ORF loss probabilities (Figure 4B). Our analyses show that asORFs are indeed protected from loss due to overlap with existing ORFs, especially when they exist in frame 1. This protection against loss increases with increasing intensity of purifying selection. Our analysis with parameters based on *D. melanogaster* was also in agreement with this result (Figure S4).

**Figure 4:**
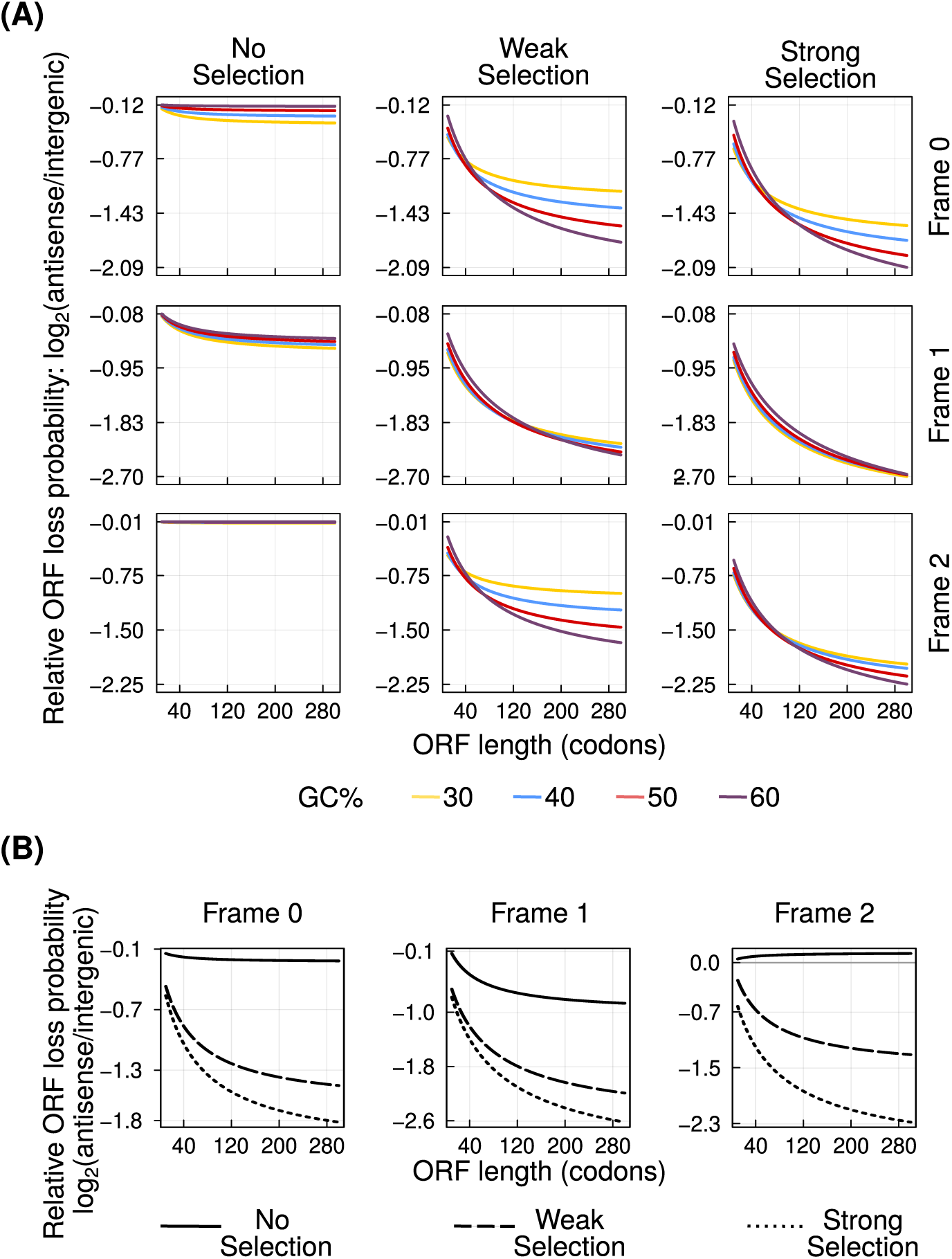
Antisense overlap can reduce ORF loss. **(A)** The probability of ORF loss in the three antisense frames (left to right) relative to that in intergenic regions (log_2_ ratio, vertical axis), at different intensities of purifying selection (top to bottom). Line colors indicate the GC-content of the ORFs. **(B)** The ORFs loss probability in the three antisense frames relative to that in intergenic regions (log_2_ ratio, vertical axis), calculated using frequencies of short DNA sequences from the yeast genome. Dotted, solid and dashed lines, denote the zero, weak and strong purifying selection, respectively. Horizontal axis in every plot shows the length of the ORFs. In every plot, we only show asORFs that overlap completely with the sense ORF.

Although antisense overlap can protect ORFs from being lost, it can also constrain the evolution of their sequence. Furthermore, effect of mutations in the sense ORF can also affect different asORFs in the three frames differently. We found that when a sense ORF is under purifying selection (weak or strong), mutational effects are the strongest for asORFs located in frame 2, and the weakest for those in frame 0 (Figure S5).

To corroborate some of our model’s predictions, we analysed the genome and the transcriptome data from the seven different lines of *D. melanogaster*. Six of these lines were obtained from different locations in Europe, whereas one line, the outgroup, was obtained from Zambia (Grandchamp *et al*., 2023). This data set allowed us to analyse gain and loss of transcripts and ORFs in short evolutionary timescales. If an asORF is found in at least one line, it is gained once in *D. melanogaster*. More specifically, the most recently emerged asORF would be detected in only one line, given the assumption that it is not independently lost in six other lines. We found that regardless of whether an asORF is present in one or many lines, they are more abundant in frame 1 than in the other two frames. This corroborates our model’s prediction that antisense overlap in frame 1 facilitates ORF gain (Figure 5A).

**Figure 5:**
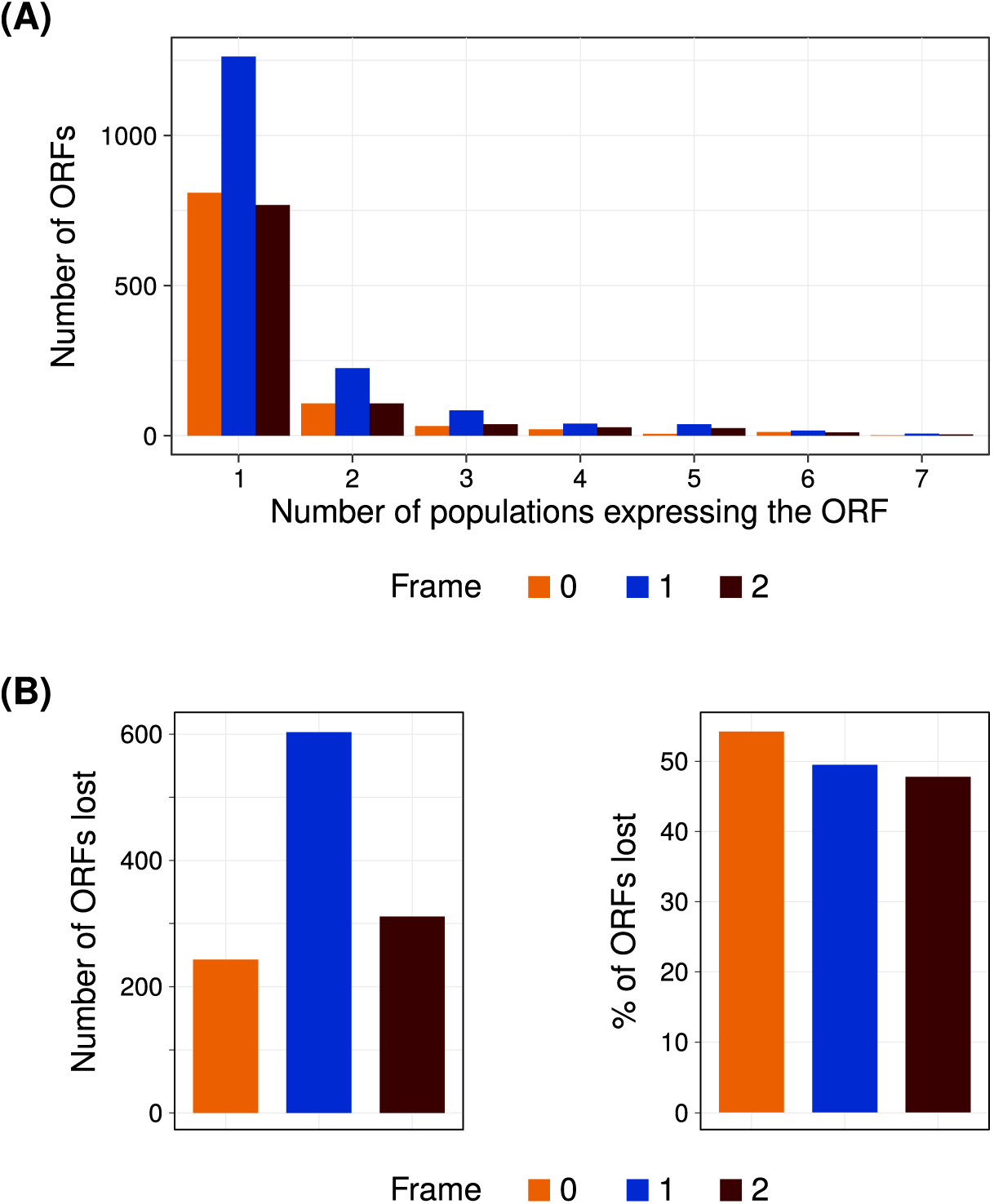
(A) Most recently gained asORFs in *D. melanogaster* are frequently located in frame 1. Horizontal axis denotes the number of *D. melanogaster* lines that contain an asORF in their transcriptome (express), and vertical axis denotes the number of such ORFs. **(B)** *D. melanogaster* asORFs in frame 0 have a higher rate of loss. First panel shows total number of lost ORFs (vertical axis) whereas the second panel shows the percentage of total asORFs that are lost. In all figure panels, the three frames are denoted by three different colors (0: orange, 1: blue, and 2: brown).

Next, we analysed the rate of ORF loss in the *D. melanogaster* lines. The genetic variance (F_ST_) between the European populations of *D. melanogaster* is low (Kapun *et al*., 2020), suggesting that they are not significantly isolated (Whitlock and McCauley, 1999). As a consequence, we could not establish a clear phylogeny for them. Thus we used a very stringent identification of ORF loss. Specifically, if an ORF is present in the outgroup line (Zambian) and at least one European line, we assume that it was lost in the rest of the European lines. For this definition, we assumed that it is unlikely for an ORF to be gained multiple times independently, and that an ORF can be shared between a European line and the outgroup only if it was already present in their common ancestor. To understand the rate of ORF loss, we normalized the number of asORFs lost in any one frame with total number of asORFs present in the same frame. We found that the rate of ORF loss was highest in frame 0, followed by frames 1 and 2 respectively (Figure 5B).

However, the magnitude of this difference was small (*<*5%) as qualitatively predicted by our model (Figure 4).

Overall, our analyses suggest that antisense overlap with an existing ORF facilitates emergence of new ORFs, and protects the existing asORFs from being lost.

## Discussion

To express a protein, a DNA sequence needs to be transcribed as well as translated. New protein coding genes can emerge *de novo* in non-genic sequences when both these requirements are met. Genomic regions that are already transcribed are thus more likely to evolve protein coding features (Iyengar and Bornberg-Bauer, 2023). Non-coding RNAs indeed harbor ORFs, and some of these ORFs are also actively transcribed, albeit less efficiently than canonical ORFs present in mRNAs (Ruiz-Orera *et al*., 2014; Ingolia *et al*., 2014; Patraquim *et al*., 2022; Wacholder *et al*., 2023). Several long non-coding RNA genes overlap with other genes in an antisense orientation (Mattick *et al*., 2023). This overlap can cause the evolution of asORFs to be constrained by the evolutionary pressures on the corresponding sense genes. The effect of ORF overlap is particularly important in viruses where novel genes frequently emerge overlapping with existing genes, in order to keep the genome compact (Sabath *et al*., 2012). In this study, we investigate how likely it is for asORFs to exist in the three possible antisense frames, and how their evolution is constrained by the purifying selection on the sense ORFs. To answer these questions, we developed a mathematical model based on mutation probabilities, and analysed the genome sequence for validating some of the model’s predictions.

Using the model, we show that asORF are most likely to be found in frame 1 than in the other two frames. This prediction is supported by our analysis of asORFs in *Saccharomyces cerevisiae* and *Drosophila melanogaster* genomes. Furthermore, asORFs in frame 1 are not only more likely to emerge, but may be also less likely to be lost than asORFs those in the other two frames. More interestingly, ORFs are generally more likely to emerge and to be found in antisense frame 1 than in intergenic regions. Conversely, these asORFs are less likely to be lost than intergenic ORFs, due to random mutations. This happens because presence of a sense ORF reduces the chances of premature stop codons to occur in the antisense frame 1.

A previous study has also investigated the effect of selection pressure on different frames, using information theory (Mir and Schober, 2014). Although this study also investigates antisense frames, its analytical approach is different from that of our model. Specifically, we calculate the probability of different kinds of mutations, and focus on the presence or absence of ORFs of different lengths, instead of measuring the fidelity of evolutionary information transfer based on relative rates of synonymous and nonsynonymous mutations. Despite these differences in the analytical approach, the findings of our study are in agreement with the previous study. That is, selection pressure on sense ORF (frame +1 in Mir and Schober, 2014) causes preservation of asORFs in frame 1 (frame −2 in Mir and Schober, 2014).

By limiting the number of tolerated mutations, an overlap with an existing ORF can affect the evolution of the protein sequence encoded in an asORF. We quantified mutational effects by estimating the average chemical difference between an original amino acid and a substituted amino acid that results due to random mutations. We found that mutational effects were the strongest in the asORFs in frame 2 (Figure S5). This means that the mutations tolerated in the sense ORFs under purifying selection produce extreme non-synonymous changes in the asORFs in frame 2.

Like all computational models, our model is based on some assumptions and simplifications, that need to be considered. For example, we use GC-content as a measure of nucleotide composition which we use in turn to calculate different probability values. For these calculations, we also use codon, dicodon and DNA trimer frequencies, which are realistic measures of nucleotide composition. Our results indeed show that probability values calculated using GC-content can sometimes noticeably differ from the values calculated using DNA oligomer distributions, especially for *D. melanogaster*. For example, our estimated probability of finding a *D. melanogaster* asORF was highest in frame 1 when we used GC-content, whereas it was highest in frame 0 when we used oligomer distributions. Both our measures of nucleotide composition can vary significantly across the genome. We used different values of GC-content for our calculations that can represent different genomic loci. In contrast, our DNA oligomer based calculations is based on the average frequency of oligomers from the whole genome. Thus they may not accurately represent any one specific locus. However, our computational framework can be adapted to analyse specific loci. Therefore, model predictions may not be 100% accurate. However, despite the possible inaccuracies, our models are able to produce results that qualitatively agree with real data. Our analysis of asORFs from budding yeast and *D. melanogaster* supports our model based finding that antisense frame 1 has higher likelihood to harbor asORFs. Our models are based on the assumptions of uniform mutation rate and independence of mutational events. These assumptions are not exactly accurate because mutation rates can vary across the genome (Monroe *et al*., 2022), and multiple nucleotides can be mutated in a single mutational event (Harris and Nielsen, 2014). Furthermore, mutation rate bias can be different in different organisms (Cano *et al*., 2022; Bergeron *et al*., 2023, also compare Table 1 and Table S1). Our results show that despite the differences in the mutation rate and mutation rate bias, between yeast and *D. melanogaster*, the results qualitatively remain the same. Thus our predictions are robust to small changes in parameters.

We believe our work opens up interesting questions, and avenues for future research. For example, the cellular functions and biochemical properties of proteins encoded by asORFs would be worth investigating. This may be especially relevant for antisense lncRNAs, some of which are involved in regulation of gene expression. asORFs may possibly provide another dimension to the cellular function of these RNAs. Translation of ORFs in lncRNAs can indeed be spatiotemporally regulated (Patraquim *et al*., 2022). asORFs may especially be relevant in organisms with compact genomes, such as viruses. Existing work indeed shows that new protein coding genes emerge in viruses, overlapping with existing genes (Sabath *et al*., 2012; Schlub and Holmes, 2020; Romerio, 2023). This overlap couples the evolution of the two overlapping genes. Eventually, understanding viral evolution may help design better therapeutic strategies against viral diseases.

## Materials and Methods

### Probabilities of finding, gaining, and losing an ORF

We calculated the probabilities of finding, gaining and losing a ORF, using nucleotide composition, mutation rate and mutation rate bias, as described in our previous study (Iyengar and Bornberg-Bauer, 2023). Briefly, a reading frame is an ORF (*P_ORF_*) when a start codon exists at its beginning (*P_ATG_*), a stop codon exists at its end (*P_stop_*), and no stop codon exists in the middle (1 *− P_stop_*). An ORF emerges (*P_ORF-gain_*) when two of the three required features are present and are not lost due to mutations, while the missing feature emerges due to mutations. Conversely, an ORF is lost (*P_ORF-loss_*) when any one of the three required features is lost. The probabilities of finding, gaining and losing an ORF containing *k* codons, are described by the following equations (Equations 1 – 3). Table 3 describes the terms used in these equations.

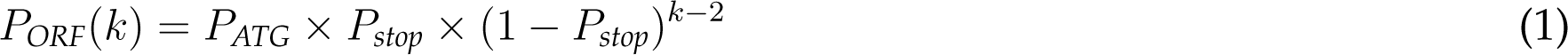

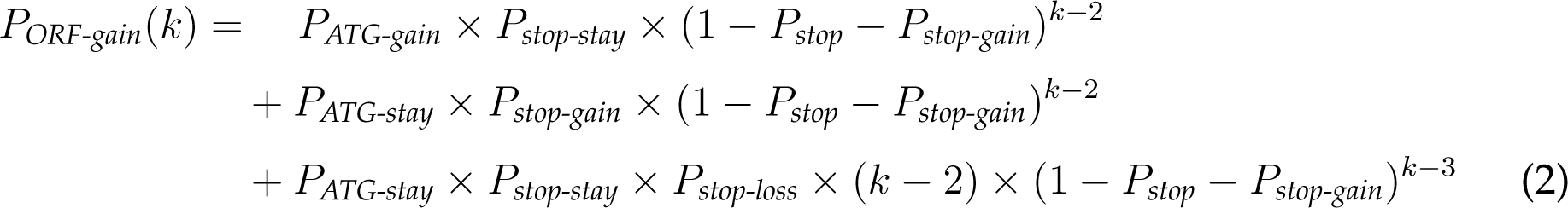

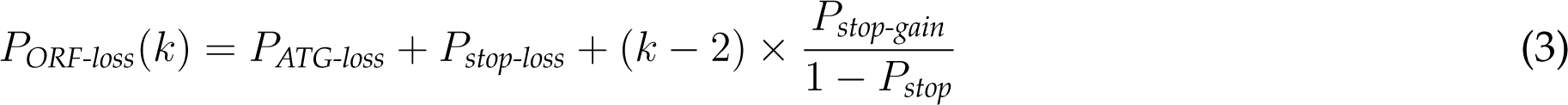

**Table 3:** Description of the probability terms used in Equations 1 – 3. Here we describe the probabilities associated with stop codons. Analogous probability terms for a start codon are denoted by the subscript, *ATG* (instead of *stop*). For asORFs, *P_stop_*, *P_stop-gain_*, *P_stop-loss_*and *P_stop-stay_*will vary depending on the frame.

| Term | Description |
| --- | --- |
| $P_{stop}$ | Probability of finding a stop codon |
| $P_{stop-gain}$ | Probability of gaining a stop codon |
| $P_{stop-loss}$ | Probability of losing a stop codon given that it already exists |
| $P_{stop-stay}$ | Probability that a stop codon exists and is not lost due to mutations |

### Modeling weak purifying selection

Both gain and loss probabilities of asORFs depend on the strength of selection on the sense ORF. That is, selection would limit the number of sense codons or dicodons that any of the existing codons and dicodons can mutate to. Under strong purifying selection only synonymous mutations are allowed, whereas weak purifying selection allows an amino acid to be substituted by a chemically similar amino acid. To determine chemically similar amino acids, we used an amino acid similarity matrix based on binding covariance of different short peptides to MHC (Major Histocompatibility Complex, Kim *et al*., 2009). As noted by Kim *et al*. (2009), we identified chemically similar amino acids from pairs of amino acids whose covariance scores are more than 0.05 (Table 4).

**Table 4:** Chemically similar amino acids identified using the data from Kim *et al*. (2009)

| Amino acid | Chemically similar amino acids | Amino acid | Chemically similar amino acids |
| --- | --- | --- | --- |
| A | P, T, V | M | I, L |
| C | - | N | - |
| D | E | P | A |
| E | D | Q | - |
| F | I, W, Y | R | H, K |
| G | - | S | T |
| H | K, R | T | A, S |
| I | F, L, M, V | V | A, I |
| K | H, R | W | F, Y |
| L | I, M | Y | F, W |

### Identification of asORFs in the genome

To identify asORFs in *Saccharomyces cerevisiae* genome, we first compiled a list of known antisense RNAs from the S288C reference genome (Engel *et al*., 2014), and combined it with the list of novel RNAs identified in a recent study (Blevins *et al*., 2021). Next, we identified all ORFs in the combined set of RNAs using the program *getorf* (Rice *et al*., 2000). Specifically, we identified the longest sequence that starts with the canonical `ATG` start codon and ends with a stop codon. We used a minimum ORF length of 30nt (default value in *getorf*). We then mapped the genomic coordinates of all the identified ORFs, verified if they overlap with a known ORF in the opposite strand, and calculated the frame of antisense overlap. We used *awk* scripts for this analysis. To calculate the number of ORFs expected from the model, we first identified genomic regions where an antisense overlap exists between an annotated ORF and a RNA. For each such region *A*, with a length *l_A_*, we calculated the total number of asORFs in each frame (nORF*_f_*) longer than 29nt (*≥* 10 codons) as:

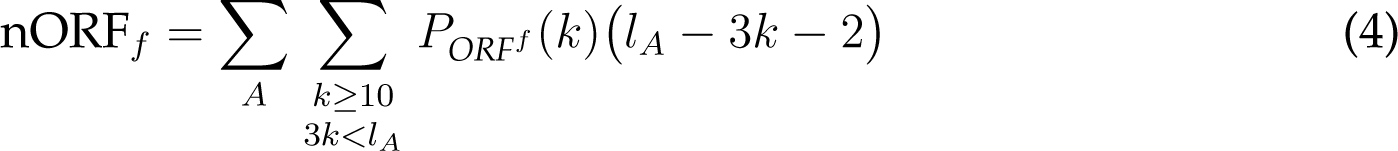

Where *P_ORF_f* (*k*) is the probability of finding an ORF in a frame *f* (Figure 1). We also identified intergenic ORFs using *getorf* (Rice *et al*., 2000).

We performed an analogous analysis for *D. melanogaster*. For details please see Supple- mentary Section 4.

### Data availability

All scripts and necessary data files are freely available on GitHub:

*BharatRaviIyengar/DeNovoEvolution*.

We implemented our model using Julia programming language using the following scripts:

- *antisenseGenes.jl* (main script)
- *antisenseGenes supplement.jl* (calculations using codon, dicodon, and intergenic trimer frequencies)
- *nucleotidefuncts.jl* (dependency for basic functions)

The *awk* scripts for asORF identification from yeast and *D. melanogaster* genome are located in the folder *DataAnalysis*. A wrapper *bash* script implements the complete analysis pipeline in both cases. We also include some original data files for yeast but not for *D. melanogaster*.

## 1. Mutation rate and mutation rate bias in in *Drosophila melanogaster*

**Table S1:**
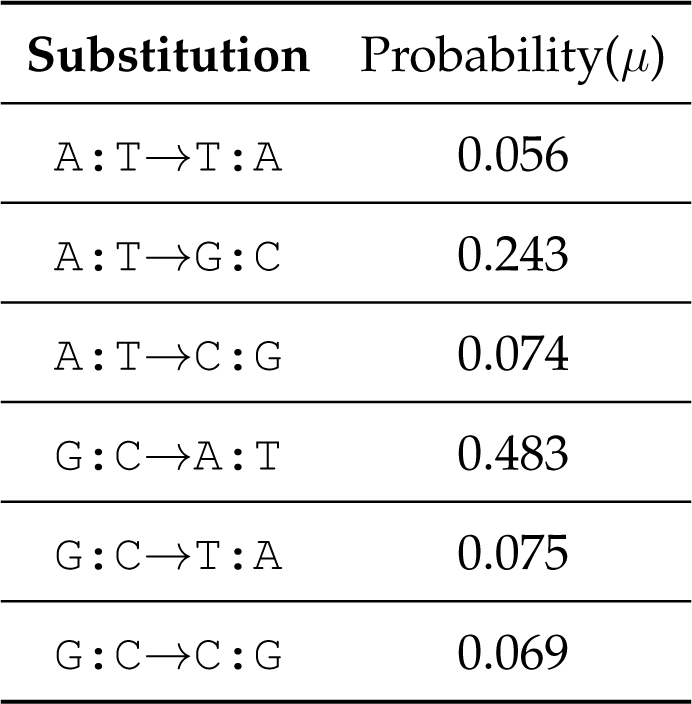
Mutation bias probabilities for different nucleotide mutations based on Schrider *et al*. (2013) and Zhang and Gerstein (2003). `A:T` denotes an `A`-`T` base pair in a double stranded DNA. Thus `A`*→*`G` mutation on one DNA strand would cause a `T`*→*`C` mutation on the complementary strand. We describe the other mutations in the same way. We used an average mutation rate of 7.8 *×* 10*^−^*^9^ mutations per nucleotide position per generation (Schrider *et al*., 2013)

| Substitution | Probability($\mu$) |
| --- | --- |
| A:T→T:A | 0.056 |
| A:T→G:C | 0.243 |
| A:T→C:G | 0.074 |
| G:C→A:T | 0.483 |
| G:C→T:A | 0.075 |
| G:C→C:G | 0.069 |

## 2. Probability of finding antisense ORFs in *Drosophila melanogaster*

**Figure S1:**
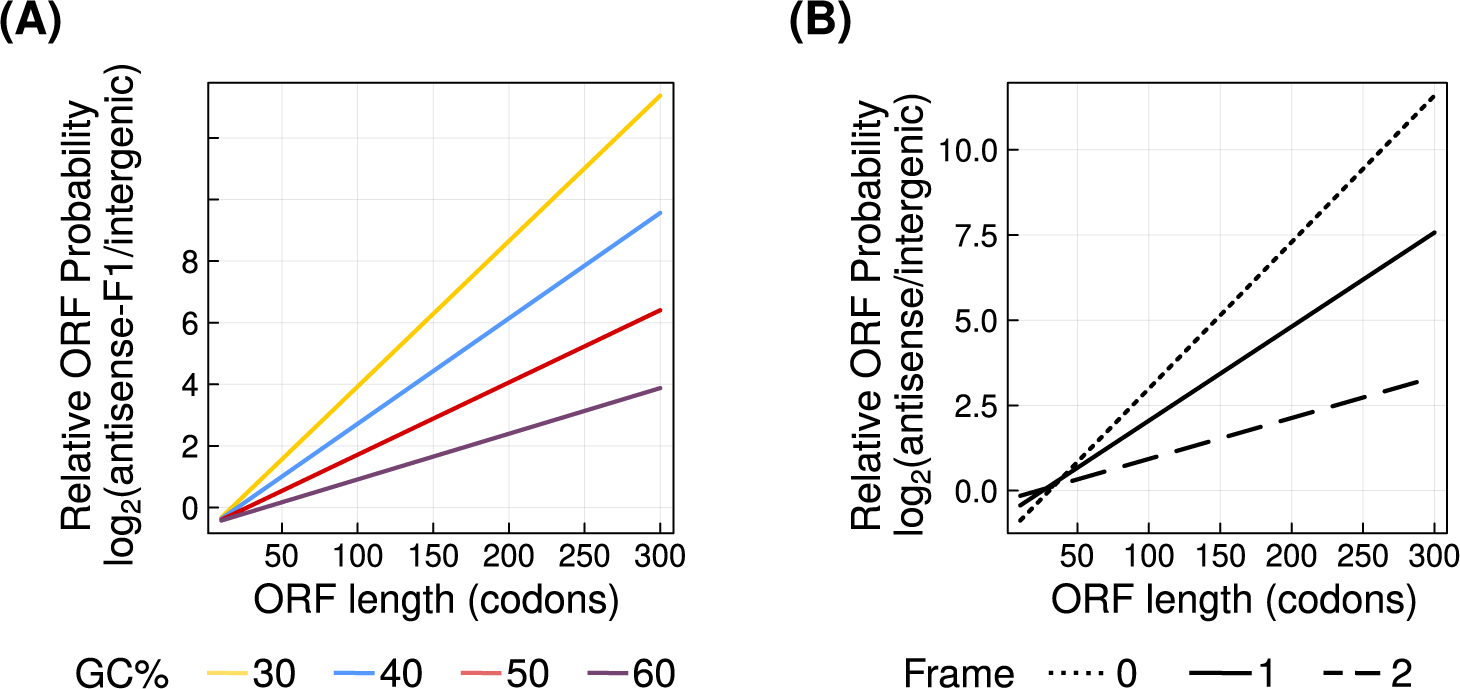
Panel **(A)** shows the probability of antisense ORFs in frame 1 relative to that of intergenic ORFs (log_2_ ratio, vertical axis). Line colors indicate the GC-content of the ORFs (yellow = 30%, blue = 40%, red = 50%, purple = 60%). We do not show antisense ORFs in frames 0 and 2 because their probabilities are identical to that of intergenic ORFs. Panel **(B)** shows the probability of antisense ORFs relative to that of intergenic ORFs (log_2_ ratio, vertical axis), calculated using frequencies of short DNA sequences from *D. melanogaster* genome. Frames 0, 1 and 2 are denoted by dotted, solid and dashed lines, respectively. Horizontal axis in both panels shows the length of the ORFs. For data in both panels, we assume that antisense ORFs overlap completely with the sense ORF.

## 3. Distribution of antisense ORFs in *Drosophila melanogaster* genome

We identified antisense ORFs and intergenic ORFs using genome and transcriptome data from seven *D. melanogaster* lines (Grandchamp *et al*., 2023a,b). We performed the same analysis for every *D. melanogaster* line. Specifically, we first identified RNAs that overlap antisense to any known protein coding gene. Next, we extracted ORFs in these antisense RNAs using *getorf* (Rice *et al*., 2000). Next, we mapped the genomic coordinates of these ORFs using nucleotide BLAST (100% query coverage and sequence identity; Altschul *et al*., 1990; Camacho *et al*., 2009), and identified all asORFs and their frame of overlap using *awk* scripts (we note that not all ORFs in antisense RNAs are asORFs). In the final step, we only analysed asORFs whose genomic sequences were uninterrupted by introns (Table S2).

**Table S2:** Summary of antisense and intergenic ORFs identified in *D. melanogaster* lines. The different asORFs reported here include sub-ORFs within longer ORFs detected by *getorf*. Here we only report asORFs that do not contain introns and that completely overlap with a protein coding exon (sense ORF).

| Line | Denmark | Finland | Spain | Sweden | Türkiye | Ukraine | Zambia |
| --- | --- | --- | --- | --- | --- | --- | --- |
| Observed igORFs | 1280190 | 1323699 | 1371327 | 1316219 | 1349122 | 1316515 | 1269809 |
| Expected igORFs | 1877477 | 1953590 | 2024171 | 1937273 | 1988489 | 1935409 | 1858742 |
| Total intergenic loci | 2147483647 | 2147483647 | 2147483647 | 2147483647 | 2147483647 | 2147483647 | 2147483647 |
| Observed asORF0 | 149 | 174 | 170 | 147 | 218 | 154 | 126 |
| Observed asORF1 | 286 | 279 | 250 | 254 | 483 | 265 | 288 |
| Observed asORF2 | 169 | 164 | 180 | 176 | 238 | 157 | 216 |
| Expected asORF0 | 123 | 123 | 133 | 126 | 189 | 126 | 126 |
| Expected asORF1 | 147 | 146 | 158 | 150 | 224 | 150 | 152 |
| Expected asORF2 | 158 | 156 | 169 | 161 | 239 | 161 | 163 |
| Total antisense loci | 589915 | 596786 | 630041 | 594764 | 928645 | 588753 | 584516 |

To estimate gain and loss of asORFs we compared their presence or absence in the transcriptome of the different *D. melanogaster* lines. We assume that an ORF emerges only once. That is, if an ORF is detected in five lines, we assume that it emerged once and spread in five lines.

As a first step, we identified ORFs that were shared by several lines. We call defined an orthogroup as a group of query unique ORF sequences detected in any of the seven lines. Our definition of orthology in this case is very stringent. For example, if an ORF duplicated in two lines, the duplicated copies belong to a different orthogroup. That is so because we were interested in the gain and loss of the original ORF and its duplicated copy separately. We also made our analysis very stringent by removing orthogroups where the ORFs from the different lines are not located in the same frame. We did so because if would be difficult to infer in which frame (line) the ORF gain occurred first.

**Figure S2:**
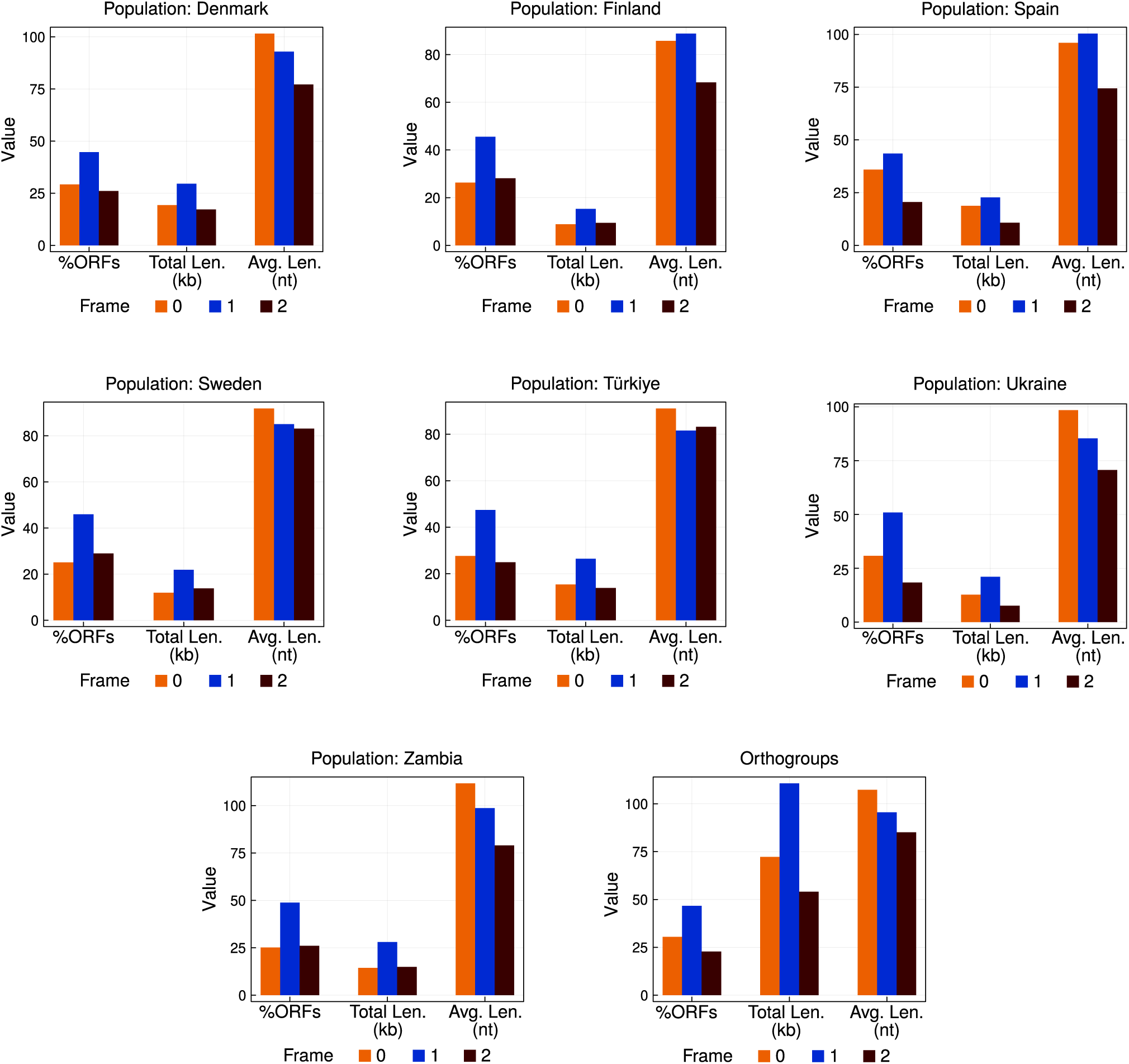
Frame preference of antisense ORFs in *D. melanogaster* genome. We show three metrics of frame preference as three bar groups – percentage of total ORFs (left), cumulative length of the overlapping region from all antisense ORFs (middle), and average ORF length (right), in each of the three frames (bar colors). We calculated these metrics from the genomics and the transcriptomics data from the seven different *D. melanogaster* lines (Grandchamp *et al*., 2023a,b).

To identify orthogroups, we used nucleotide BLAST(Altschul *et al*., 1990; Camacho *et al*., 2009) and restricted alignments to those with a high score and 100% coverage. We also ensured that all homologous ORFs were antisense-overlapping with the same established gene. Most orthogroups contained only one ORF per line. However, some orthogroups contained several ORFs in a single line, due to tandem duplications. We split these orthogroups such that they contained only one ORF per line, and sorted them according to their frame and relative genomic position. Among the 3536 orthogroups we detected, 105 had several ORFs in several lines. 32 out of these 105 orthogroups contained more than four duplicates in some lines. We discarded these orthogroups because we could not reliably categorize them into sub-orthogroups after splitting them based on frame and position. We also discarded 147 or-thogroups were from our analysis because the homologous ORFs were located in different frames.

To estimate the loss, we used the outgroup (Zambian) line. If an ORF was found in the out- group and at least one European line, we assume that it emerged in an ancestral *D. melanogaster* population and was lost in rest of the five European lines. We found 319 orthogroups where the ORF was present in the Zambian line and at least one European line but not all six of them.

## 4. Gain and loss probabilities of antisense ORFs in *Drosophila melanogaster*

**Figure S3:**
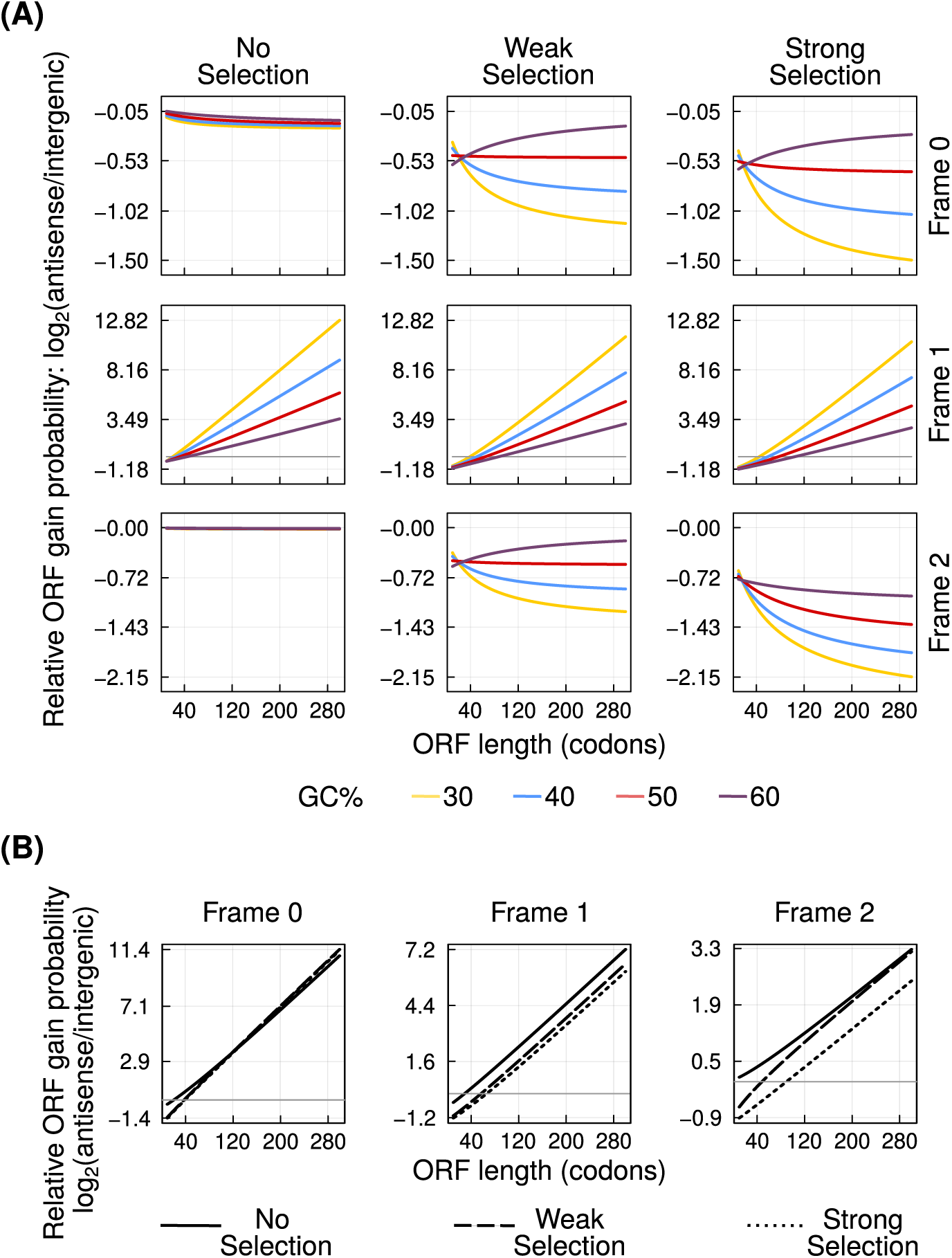
Antisense overlap can facilitate ORF emergence. Panel **(A)** shows the probability of ORF emergence in the three antisense frames (left to right) relative to that in intergenic regions (log_2_ ratio, vertical axis), at different intensities of purifying selection (top to bottom). Line colors indicate the GC-content of the ORFs. Panel **(B)** shows the ORFs gain probability in the three antisense frames relative to that in intergenic regions (log_2_ ratio, vertical axis), calculated using frequencies of short DNA sequences from *D. melanogaster* genome. Dotted, solid and dashed lines, denote the zero, weak and strong purifying selection, respectively. Horizontal axis in all panels shows the length of the ORFs. For data in both panels, we assume that antisense ORFs overlap completely with the sense ORF.

**Figure S4:**
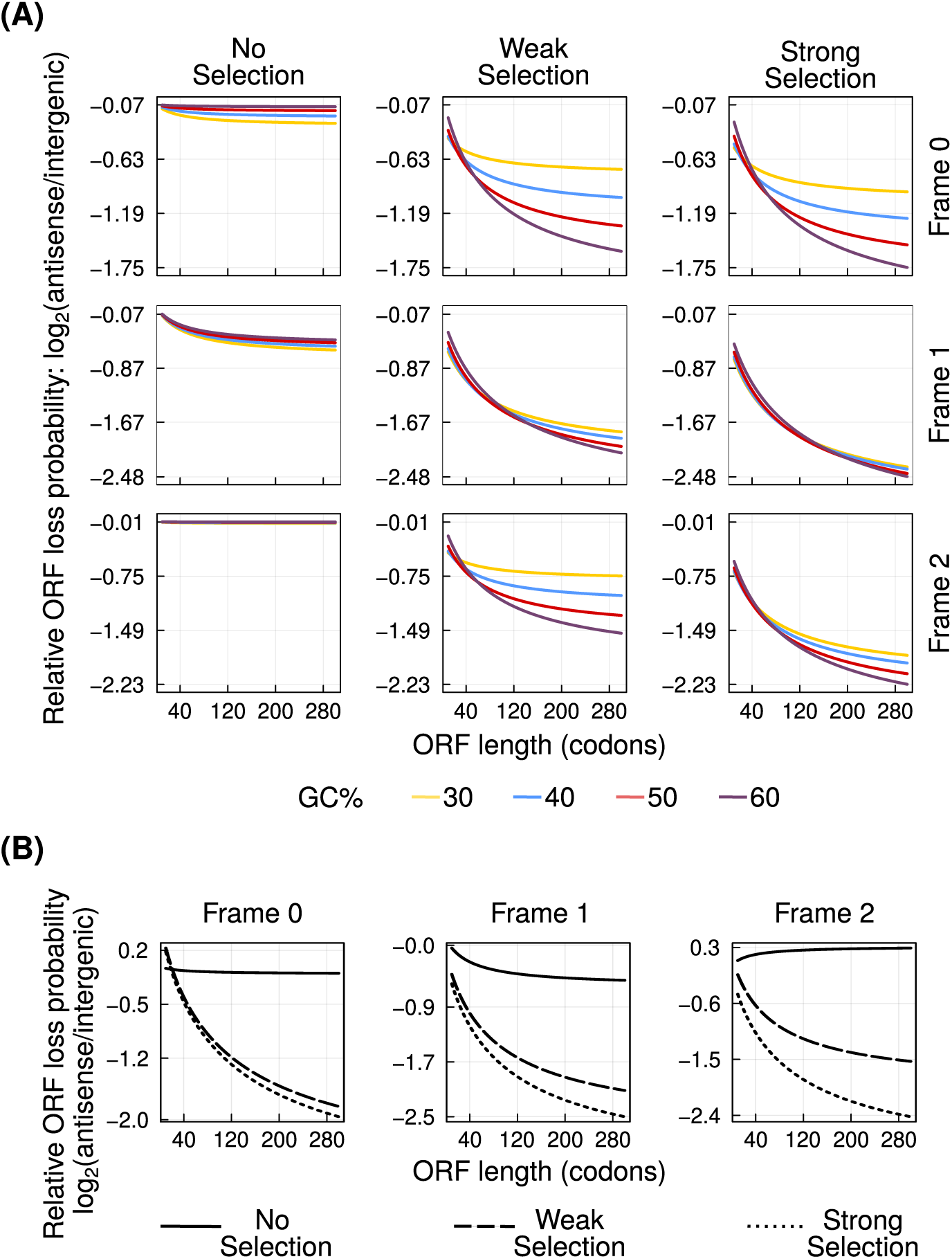
Antisense overlap can reduce ORF loss. Panel **(A)** shows the probability of ORF loss in the three antisense frames (left to right) relative to that in intergenic regions (log_2_ ratio, vertical axis), at different intensities of purifying selection (top to bottom). Line colors indicate the GC-content of the ORFs. Panel **(B)** shows the ORFs loss probability in the three antisense frames relative to that in intergenic regions (log_2_ ratio, vertical axis), calculated using frequencies of short DNA sequences from *D. melanogaster* genome. Dotted, solid and dashed lines, denote the zero, weak and strong purifying selection, respectively. Horizontal axis in all panels shows the length of the ORFs. For data in both panels, we assume that antisense ORFs overlap completely with the sense ORF.

## 5. Effect of mutations on asORFs

In the previous sections, we showed that purifying selection on the sense ORF can affect the emergence and loss of asORFs. We next asked if this purifying selection can also constrain the diversification of the proteins encoded by asORF sequences. To this end, we first calculated the “chemical distance” (*δ*) between any two amino acids. For this calculation we used a distance matrix that we derived from an experimentally estimated amino acid similarity matrix reported in a previous study (Kim *et al*., 2009). Next, we calculated the average chemical difference (*δ̄*) introduced by a random mutation, weighted by the probability of different mutations (Equation 1). To this end, we created an amino acid distance matrix by modifying the amino acid similarity matrix of Kim *et al*. (2009). Specifically, we subtracted the value of 0.3 from each element of the matrix, reversed the sign of each element, and set the diagonal to zero. By doing this, we set every distance value to be greater than 0. Next, we calculated the average chemical difference introduced by any mutation (*i → j*) allowed under a selection regime. Specifically, if *i* denotes the original codon, *j* denotes the substituted codon, *P_i_*denotes the probability of finding codon-*i*, *µ_ij_* denotes the probability of codon-*i* mutating to codon-*j*, and *δ_ij_* denotes the chemical difference between the amino acids encoded by these codons, then the average chemical difference is defined by the following equation:

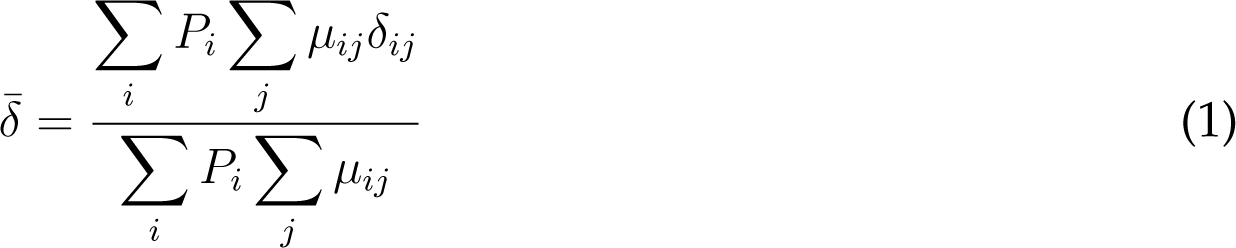

Using *δ̄* as a measure of divergence, we estimated the extent to which asORFs in the three frames can diverge as a result of mutations, and due to purifying selection on the sense ORFs. Likewise, we also calculated the divergence of intergenic ORFs as a consequence of random mutations. We found that frame 2 allows maximum divergence of asORFs, under both weak and strong purifying selection on the sense ORF (Figure S5A). asORFs in frame 0 diverge the least. Interestingly, strong selection on sense ORFs increases the divergence of asORFs in frame 2. The reason could be that the few mutations that do occur under strong purifying selection, cause a relatively higher increase in divergence than the more numerous mutations that are allowed to occur under weak purifying selection. We also found that the divergence of asORFs in frame 2 was higher than than that of intergenic ORFs under both selection regimes. We note this result does not mean that intergenic ORFs can diverge less than asORFs. Evolution of intergenic ORFs is not constrained by another DNA sequence. However, as long as the mutants do not affect the organismal fitness, evolution would not be biased towards divergence increasing mutations. Thus random mutations in intergenic ORFs could also consist of many synonymous and chemistry preserving mutations, that are probably disallowed in frame 2 asORFs due to purifying selection on sense ORFs.

In contrast to frame 2, the divergence of asORFs in the other two frames decreased with increasing strength of purifying selection on the sense ORF (Figure S5A). For example, asORFs in frame 0, did not diverge at all when the sense ORF was under strong purifying selection. asORFs in frames 0 and 1 also diverged less than intergenic ORFs under both selection regimes.

We observed identical trends in divergence of asORFs from our analysis based on *D. melanogaster* parameters (Figure S5B).

These findings not negate the fact that intergenic ORFs have less constraints on their evolution. Even though chemical consequences of tolerated mutations may be larger in some asORFs than in intergenic ORFs, purifying selection on the sense ORF limits the total number of possible mutations. This would not be the case for intergenic ORFs.

**Figure S5:**
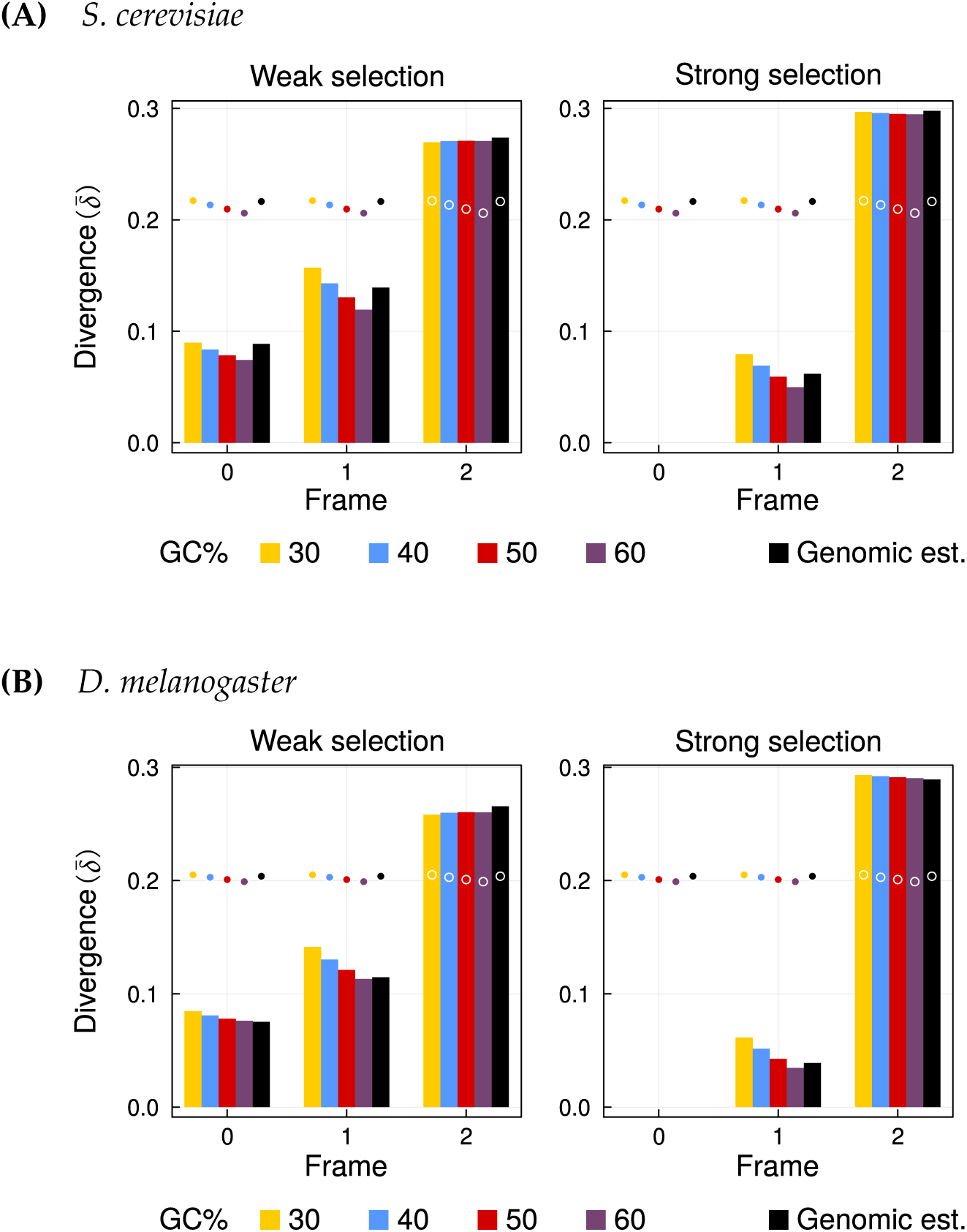
Antisense ORFs in **(A)** *S. cerevisiae* and **(B)** *D. melanogaster*, can diversify when sense ORFs are under purifying selection. Vertical axis denotes the divergence (*δ̄*) of asORFs due to a random mutation when the sense ORF is under weak (left) or strong (right) purifying selection. Horizontal axes denote the three antisense frames. Colored bars denote divergence values of asORFs with different GC-content, and black bars denote the diversity values calculated using frequencies of short DNA sequences from the yeast genome. Filled circles that are similarly color coded, denote the divergence of intergenic ORFs due to mutations.

